# Meta-PepView: a metaproteomics performance evaluation and visualization platform

**DOI:** 10.64898/2025.12.03.692192

**Authors:** Ramon van der Zwaan, Berdien van Olst, Mark C.M. van Loosdrecht, Martin Pabst

## Abstract

Microbial community proteomics is rapidly gaining traction as it allows exploration of functional processes in microbial ecosystems. Consequently, there is a growing need for user-friendly tools that enable performance evaluation and interactive visualization of the increasingly complex community proteomics data. We introduce meta-PepView, a web-based platform that enables performance evaluation and interactive visualization of metaproteomics data, ensuring transparent and reproducible metaproteomics experiments. Meta-PepView integrates spectral sequencing outputs and databases from common proteomics search engines and classification tools. It runs efficiently on laptop or desktop PCs and can be deployed as a Docker container or installed with pip, from which it is operated through the web browser. Its modular design allows easy expansion with new data sources and annotation databases. Meta-PepView is an open-source Python platform that is freely available under the Apache License 2.0. The code is available on GitHub: https://github.com/ramonzwaan/metapepview/. Here, we showcase the meta-PepView platform with data from synthetic communities and previously published microbiome studies.

## INTRODUCTION

Metaproteomics has emerged as an essential component of the meta-omics toolbox, as it allows to explore the expressed functions and interactions within microbial communities (1-4). Consequently, metaproteomics is now widely applied to study natural ecosystems such as soils (5-8), oceans (9-11), the human microbiome (12-16), and engineered ecosystems including processes in wastewater treatment and biogas plants (17-21). In addition, metaproteomics supports the development of resource recovery strategies, including the recovery of valuable biopolymers from microbial biomass (22-24) and the discovery of novel enzymes (25).

Compared to conventional single species proteomics, metaproteomics faces additional challenges, including difficult sampling from natural environments, increased sample matrix complexity, protein extraction from microbes with diverse cell physiologies, and often high microbial diversity and unevenness. Consequently, metaproteomic experiments require careful experimental implementation and data processing that account for taxonomic complexity and reference databases covering the entire microbiome (26-31). However, despite recent advancements in metaproteomics (32, 33), the accuracy and reproducibility can be constrained by poor experimental quality control and limited method standardization across laboratories (4, 34).

The need for reproducible experimental procedures is widely recognized, as the scientific community has already reported a “reproducibility crisis” (35), highlighting the growing difficulty of reproducing published data (34). Recent inter-laboratory studies in the omics field have tested reproducibility and comparability, often with sobering results, as individual labs frequently produce widely varying outcomes (4, 9, 36, 37). While much of this variability stems from differing methods, a considerable portion presumably also arises from suboptimal application and inadequate performance control of the techniques used (34). This is further compounded by the lack of tools that allow experimental evaluation and straightforward visual interpretation of metaproteomics experiments.

Several bioinformatics tools have been developed to support the analysis and visual interpretation of metaproteomics data, including MetaProteomeAnalyzer (38), which allows spectral data processing and visual metaproteomics analysis within one application, MetaQuantome (39), which provides a pipeline for quantification and cross-sample fold change analysis of taxa and functions via a command line interface, and Prophane (40) and MetaGOmics (41), which offer taxonomic (Prophane only) and functional classification of metaproteomics data via a web-based interface. Both latter tools operate on a job-based manner where input data are imported into the interface and processing is performed in the background. In addition, the Unipept web services, (42) and the recently released PathwayPilot (43) provide a useful resource to annotate and visualize taxonomies and functions based on peptide sequences matched to the UniprotKB reference database. Furthermore, Galaxy-P (44) offers an extensive modular online multi-omics data processing platform that allows integration of a wide collection of processing tools (including MetaQuantome-based tools). While these tools meet a wide range of user needs, interactive visualization to support community analysis and metaproteomic performance evaluation is not directly provided.

The need of performance evaluation and quality control tools within proteomics workflows has been recognized since the early stages of proteomics (45-51). Recently, the Human Proteome Organisation also recommended standardising reporting of quality parameters using the mzQC format (47). Common performance parameters for proteomics experiments include an effective chromatographic separation leading to an even distribution of peptide precursors across the separation gradient (52), a high peptide precursor fragmentation rate, a good fragment ion coverage (53), a high peptide precursor transmission, a comprehensive coverage of the employed reference database (54, 55), and the depth of taxonomic and functional classification for the identified peptides or proteins (56). Nevertheless, although existing tools have become vital for the metaproteomics community, a user-friendly platform that combines metaproteomic performance evaluation with interactive community data exploration has so far been lacking.

Here, we introduce meta-PepView, a user-friendly web interface supporting visualization and performance evaluation of metaproteomics data. Meta-PepView integrates data from database search outputs including MaxQuant (57), PEAKS (58), and Sage (59, 60), taxonomic classifications based on NCBI (61) and GTDB (62), and functional annotations from eggNOG-mapper (63, 64), KEGG orthology classifications or GhostKOALA (65). Functional mapping uses the KEGG pathway visualisation resource (66). The performance evaluation module extracts parameters from mzML and featureXML files, *de novo* and database search files. It provides insights into parameters such as precursor density, fragment ion coverage, precursor transmission, reference sequence database coverage, taxonomic lineage drop-off including benchmarking to large reference datasets. The lightweight open-source platform can be installed on any laptop or desktop via Docker or pip and runs in standard web browsers. Its modular design allows easy addition of new data sources and databases. Meta-PepView is freely available on GitHub: https://github.com/ramonzwaan/metapepview/.

### EXPERIMENTAL PROCEDURES

#### Meta-PepView

documentation about installation and use of meta-PepView is found at https://ramonzwaan.github.io/metapepview/. The open-source Python code and docker files are freely available via GitHub: https://github.com/ramonzwaan/metapepview/.

##### Dashboard architecture

Meta-PepView is presented as a web interface that can be set up locally on any PC platform and accessed through the web browser. The tool is developed in Python using the Dash framework as graphical interface and the Plotly graph library for the dashboard plots. Set up of the dashboard server is currently facilitated via a Docker image.

##### Data input formats

For microbial community visualization, the tool supports metaproteomics data from various ‘database search’ and ‘*de novo*’ peptide identification engines: Peaks (database search, *de novo*) (58), MaxQuant (database search) (57), Sage (database search) (59), Novor (*de novo*) (67), and Casanovo (*de novo*) (68). It processes peptide identification data described at the spectral match level: each peptide spectrum match is described separately without aggregation of peptide sequences or protein groups. To classify peptides to taxonomy and protein function information, meta-PepView supports import of taxonomy and function mappings. For taxonomy, it expects a plain text tabular file format that maps protein ID or peptide sequence (present in database search data) to taxonomy ID from NCBI (61) or GTDB (69). Alternatively, taxonomy output from KEGG GhostKOALA (65) may be provided. For function classification, standard output files from eggNOG-mapper (63) or KEGG GhostKOALA (65) are supported. For experimental quality evaluation, meta-PepView supports spectral data import in mzML format in conjunction with feature data in featureXML (generated from OpenMS (70)) format, as well as database search input and *de novo* input data in any format as described above.

##### Importing online reference databases

For taxonomy and functional classification of peptides, meta-PepView consults public databases available online: the NCBI taxonomy database (71), the GTDB taxonomy database (69), and a KEGG mapping dataset from KEGG orthology IDs to pathway and BRITE groups (72). Download of latest datasets is managed through the dashboard interface and is required during initial setup of the tool.

##### Determining community composition

Meta-PepView can store several samples inside a multi-sample dataset (the project table). Each sample contains peptide identification information grouped by sequence (match count, integrated signal value, confidence, *etc*.), with corresponding taxonomy and function annotation. A new sample can be added to the table by importing metaproteomics data together with taxonomy and function information. Meta-PepView will process the data and append it as a new sample to the project table. Before sample import, consensus of input sources (identification engine, taxonomy and function database, etc.) between the new sample and the present project table is asserted to ensure consistency. During import, database search and *de novo* peptide identifications are separately grouped by peptide sequence. Peptides are quantified by identification count and summed signal intensities (if present in the input data). Next, peptide sequences are supplemented with taxonomy and functional information from the supplied import files. In case of multiple taxonomy ID assignments for a single peptide sequence, the last common ancestor (LCA) is stored for the peptide. For multiple function assignments, consensus for function ID is checked and in case of conflicts, either no ID is assigned, or the IDs are concatenated depending on user selection. Optionally, peptide sequence may be taxonomically classified by matching against a public sequence database. Here, meta-PepView classifies peptides with taxonomies from UniprotKB through the Unipept API (73).

##### Evaluating experimental performance

Meta-PepView supports visualization of experimental performance parameters from raw MS spectral files in mzML format. The current visualization module was designed and tested with Data-Dependent Acquisition (DDA) data as obtained from quadrupole Orbitrap mass spectrometers (Thermo Fisher Scientific). MzML files can be generated from raw spectral files with MSConvert (74), using zlib compression and peak picking at MS1 and MS2 level. Feature information can be added into the module by importing a feature map in featureXML format, which can be generated by the “FeatureFinderCentroided” tool from OpenMS (70). In addition, the spectral data can be supplemented with database search and *de novo* identification data to include identification performance to the analysis. Here, peptide properties and confidences are matched to their corresponding spectral IDs. To ensure correct matching of datasets, source file (raw spectral file) names are checked between spectral and peptide data. Meta-PepView performs “benchmarking” using shotgun proteomics datasets obtained from ∼100 yeast experiments (*S. cerevisiae* obtained from either an in house collection (75) or Promega Cat. No. V7461) analysed on a Q Exactive Plus Orbitrap mass spectrometer and 12 CAMPI SIHUMIx experiments (S01-S12) (4). Raw data were database searched and *de novo* sequenced using Peaks Studio X, employing either the *S. cerevisiae* reference proteome or the SIHUMIx metagenome. For user generated benchmark datasets, a command line tool to build summary files (“mpv-buildref”) is supplied that processes the experiment dataset into a summary file for import.

##### Used demo datasets

Three published datasets were analysed in the meta-PepView dashboard. Raw spectral data of four biological replicates from the Kleiner “Equal protein” mock community (Run 2 P1– 4) (76) were downloaded from the Proteomics IDEntifications Database (PRIDE) (PXD006118) and were analysed with database search matching against the “Mock_Comm_RefDB_V3.fasta” database. Additionally, raw spectral data from SIHUMIx communities used in the CAMPI study (4) were downloaded from PRIDE (PXD023217) and were analysed (S01-S12, 15 raw files in total) with database search matching against the “SIHUMI_DB2MG.faa” database and “SIHUMI_DB1UNIPROT.faa” database. Finally, raw spectral data from two replicates from a *Clostridium* enrichment culture were downloaded from PRIDE (PXD040972) and were analysed (77) with database search matching against the “Clost_K_MG_DB.fasta” database. Database search and *de novo* sequencing from raw spectral datasets were processed with Peaks Studio 10 (58). For the “equal protein” mock community and SIHUMIx community data, the following analysis settings were set for database Search and *de novo* sequencing: Parent ion charges between 2–8 were considered. The parent mass error tolerance was set to 20 ppm, while the fragment mass error tolerance was set to 0.02 Da. A maximum of 2 missed cleavages was allowed for the peptides. For modifications, Carbamidomethylation was set as fixed modification and variable modifications of Deamidation (NQ), Oxidation (M), and Acetylation (N-term) were set. A maximum of 1 variable modification per peptide was set. For database search reporting, database search peptide-spectrum matches (psms) were exported at a false discovery rate (FDR) threshold of 1%. For *de novo* reporting, *de novo* sequences were exported at a ALC threshold of 50%. For the *Clostridium* enrichment samples, database search and *de novo* datasets were taken from the NovoLign study (27). Raw spectral data were converted to mzML format with MSConvert (78), using the Peak Picking filter for MS1 and MS2 scans. The protein databases used for database search matching were taxonomically annotated and converted into a protein ID – taxonomy ID mapping dataset for import into meta-PepView. For the “equal protein” database by Kleiner, protein ID and taxonomy information was extracted from each protein fasta header and converted to NCBI taxonomy ID representation (SI EXCEL DOC). For SIHUMIx and *Clostridium* enrichment samples, the “SIHUMI_DB2MG.faa” (SIHUMIx) and “Clost_K_MG_DB.fasta” (*Clostridium*) protein databases were taxonomically classified by sequence alignment against the Uniref100 database (Downloaded on 9-5-2025) (79) with Diamond (80). Diamond was operated with the following settings: --fast, --algo 0, --matrix BLOSUM62, --top 5, --evalue 0.0005. NCBI taxonomy IDs were extracted from the matched Uniref100 protein headers. For proteins with multiple taxonomy ID matches, the LCA was determined by calculating the “bit LCA” (27). Functional annotation of the protein sequence databases was done with eggNOG-mapper (63). EggNOG-mapper was run in the default configuration. For SIHUMIx samples database search matched with the “SIHUMI_DB1UNIPROT.faa” database, taxonomic annotation was performed in the “peptide-centric” method as described in the CAMPI study (4), using Unipept to match peptide sequence to taxa, and filtering all taxa not part of the SIHUMIx community out prior to LCA determination.

## RESULTS

### Overview of the dashboard interface

The meta-PepView (v0.1) dashboard interface can be operated through a web browser by navigating to http://localhost:8050 after starting the docker server. The individual modules are accessible from the sidebar menu, which contains the main sections: i) Project management, ii) Community analysis, iii) Experimental evaluation and iv) Online resources. Each section contains one or more modules. The functions of each module are summarized in Figure 1, while an illustration of each module interface is provided in Figure 2. The sidebar allows users to download public taxonomic and functional databases, and once successfully loaded, the status indicator turns green.

**Figure 1.**
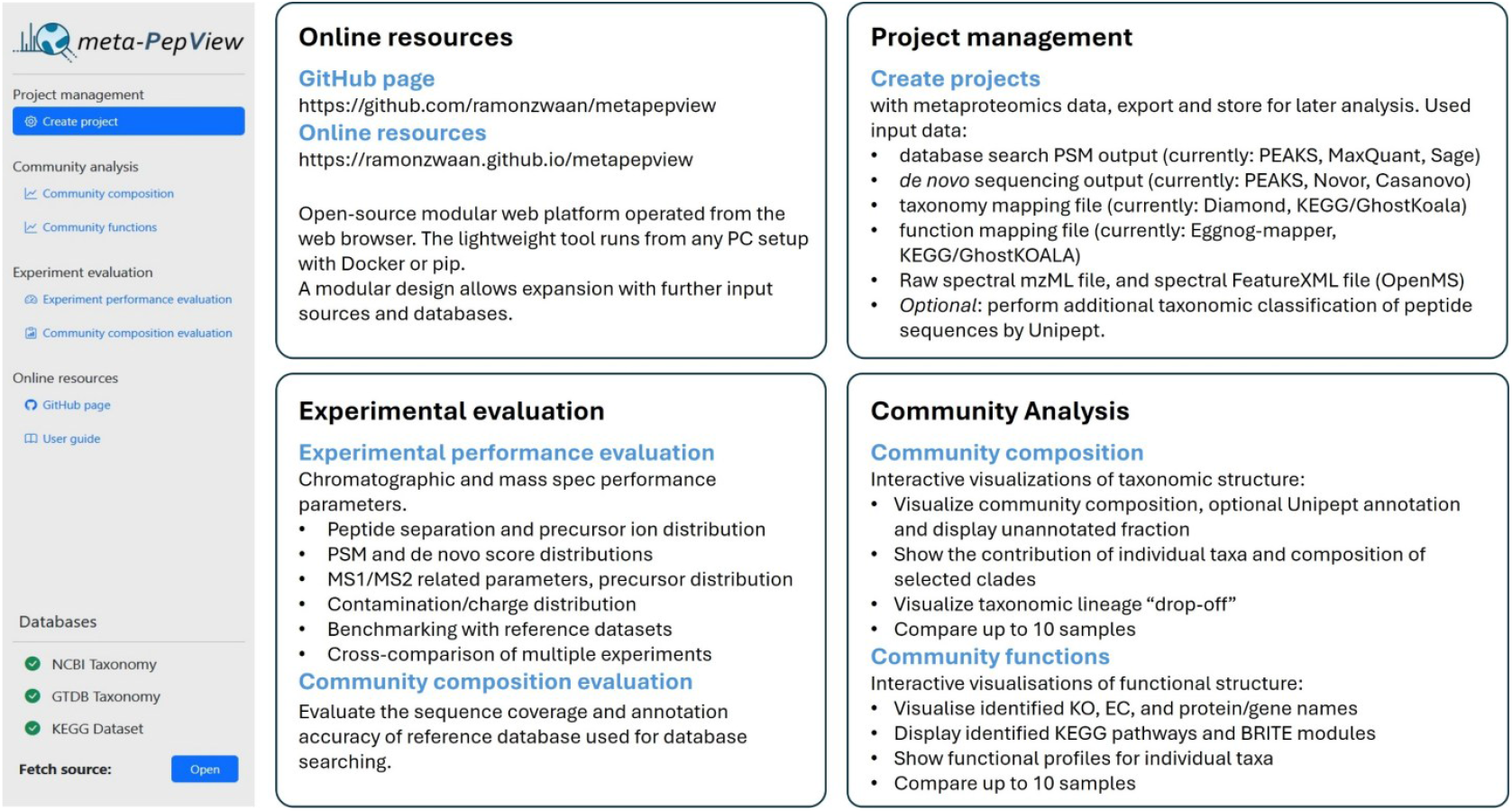
Overview of meta-PepView modules and functions (version 03/12/2025). The meta-PepView sidebar contains three main modules: Project management (“Create project”), Community analysis (“Community composition”, “Community functions”), and Experimental evaluation (“Experiment performance evaluation”, “Community composition evaluation”). The “Create project” module manages the import of metaproteomics experiments into meta-PepView. The “Community composition” module provides interactive taxonomic composition graphs, *heatmaps and the* taxonomic lineage drop-off. The “Community functions” module displays KEGG profile bar graphs *and* pathway maps. The “Experiment performance evaluation” module *provides* performance *evaluation parameters, allows* comparison of multiple samples, and benchmarking against reference datasets. Finally, the “Community composition evaluation” module assesses the reference database used for database searching in terms of sequence coverage and taxonomic classification. “Fetch source” downloads the most recent versions of the taxonomy nodes, whereas the “Databases” panel indicates which databases are currently loaded.

**Figure 2.**
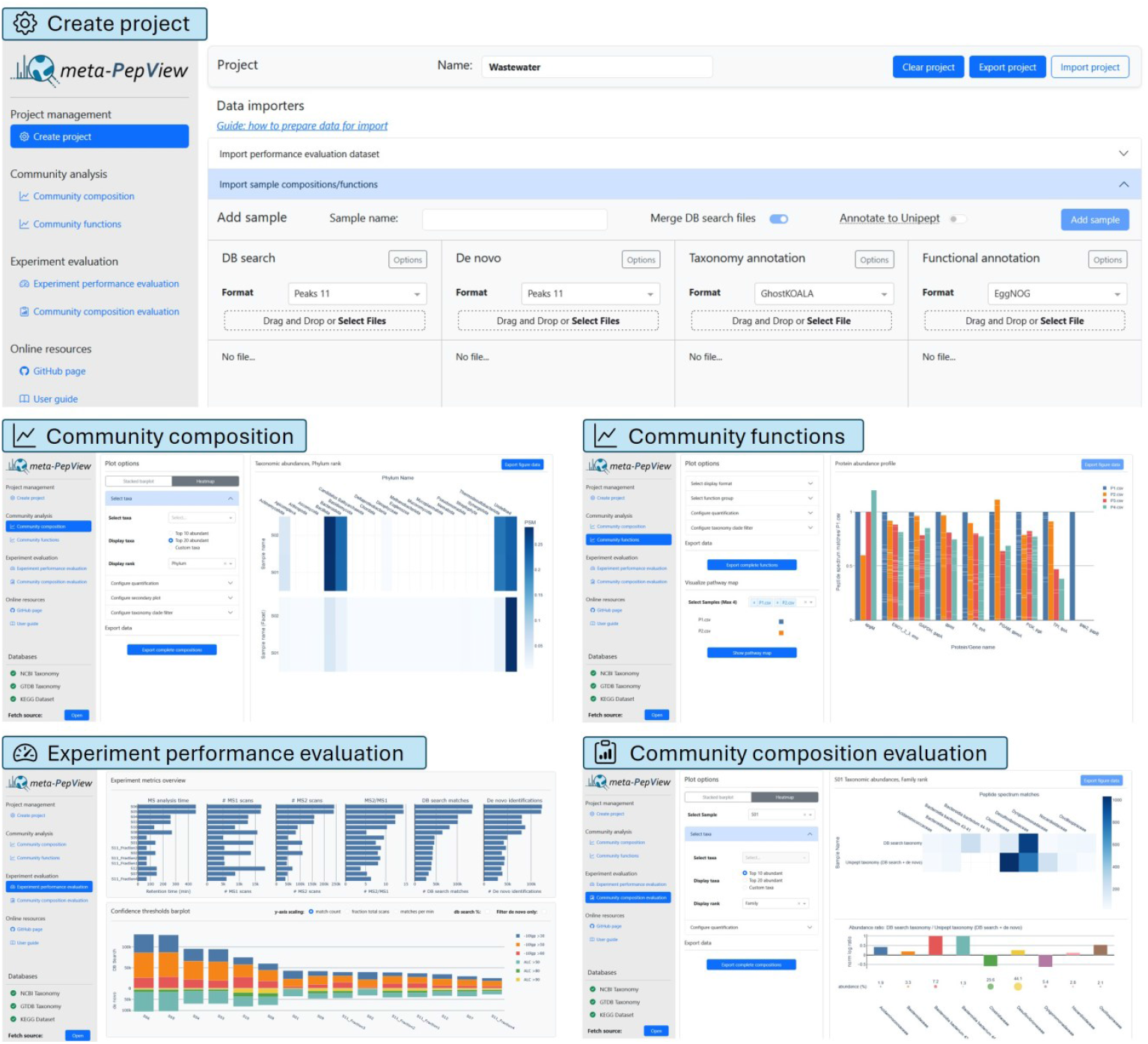
Illustration of meta-PepView (v0.1) module interfaces: Create management (Project Management), Experimental performance evaluation (Experimental evaluation), Community composition evaluation (Experimental evaluation), Community functions, and Community composition (Community analysis).

The “Create project” module allows import and export of “projects” into meta-PepView. Here, metaproteomics samples are loaded for visualization in the community analysis and experimental evaluation modules. For visualization in the community analysis module, meta-PepView supports import of multiple samples. Each sample requires database searching outputs from either MaxQuant, PEAKS or Sage, a protein ID-taxonomy file, such as obtained from Diamond (80) or GhostKOALA (65), and a protein ID-function file from either eggnog-mapper (63, 64) or GhostKOALA (65). Optionally, *de novo* sequencing data from PEAKS or Novor (67, 81) can be included. Additional taxonomic and functional annotation of the peptides can be performed by meta-PepView using the NovoBridge workflow through the Unipept database (26, 42), by selecting “Annotate to Unipept” in the “Add sample” window. A project can also store one additional sample for evaluation in the performance evaluation module. This sample requires a raw mzML spectral file, database searching outputs from either MaxQuant, PEAKS or Sage, and *de novo* sequencing data from PEAKS or Novor (67, 81). Advanced evaluation of the chromatographic separation requires an additional OpenMS feature file (featureXML) (70). Currently, meta-PepView has been tested with mzML files generated from Thermo Orbitrap systems in data-Dependent acquisition (DDA) mode. For more information on format support and preparation of input files for import, see the meta-PepView documentation: https://ramonzwaan.github.io/metapepview/ (82).

The “Community composition” module provides taxonomic abundance bar graphs and heat maps for different taxonomic ranks. This can be done with different quantifiers (psms, areas) and side by side with graphs adjusted for the “unannotated” fraction, additional Unipept sequence annotation, among other parameters.

In addition, this module provides the taxonomic lineage “drop-off”, referring to the degree of peptide uniqueness across the taxonomic lineage (Figure 4 A–B). The “Community function” module visualizes the detected KEGG elements (KO, EC, module, and gene names) as bar graphs and as interactive Kegg pathway maps (66).

The “Experiment evaluation” module contains the “Experiment performance evaluation” and “Community composition evaluation” modules. The “Experiment performance evaluation” module determines a range of quality parameters, including MS1/MS2 scan ratios, ion transmission efficiency, elution densities and peak widths, contamination index, score distributions, etc., which are linked to metaproteomic performance. The module also supports direct assessment of a large set of experiments. For this, meta-PepView provides the command line tool *mpv-buildref*, which generates quality-parameter summary files from metaproteomics experiments (82). These can be imported into meta-PepView and enable benchmarking of individual metaproteomics experiments against reference datasets, or cross-comparison of multiple experiments. SIHUMIx and yeast reference datasets are included automatically with the installation. The “Community composition evaluation” extends the “Community composition” module and provides additional assessment of sequence coverage and the taxonomic classification of the protein reference database used for database searching. It compares the taxonomic profiles obtained from database search peptides and local taxonomy mapping with those derived from *de novo* (and optionally database search) sequences annotated via the Unipept database. Details on installation and operation of meta-PepView can be found in the online documentation: https://ramonzwaan.github.io/metapepview/ (82). In the following sections, we demonstrate the application of meta-PepView to metaproteomics experiments on synthetic and natural communities described in previous studies.

### Example A: visualisation and analysis of community composition and functions

In the following section, we demonstrate the MetaPepView “Community analysis” section (“Community composition” and “Community function” modules) by exploring the synthetic Kleiner “equal protein community” (76) and the SIHUMIx mock community (83). The Kleiner “equal protein community” was created by combining 20 known species (23 strains) such that their protein quantities are equal (4.26 % per strain). Additionally, 5 phages were added at 10x lower protein quantity compared to the other microbes (0.426 % per phage). Some of these strains are closely related, challenging the analysis. A project with four replicate metaproteomics experiments (“run 2” P1–4) (76) with corresponding PEAKS database search outputs (26, 27), de novo sequencing, and taxonomic and functional classification files were imported into meta-PepView. An additional taxonomy mapping file was imported, generated from the provided taxonomy data (76), and a function mapping file, as obtained from EggNOG-mapper. The taxonomic compositions between the replicates were analysed in the “Community composition” module (Figure 3A). An interactive taxonomic bar plot of the species compositions for each replicate, alongside the expected composition based on the known protein quantities, is shown in Figure 3B. At first glance, the four replicates largely show the expected even distribution of community members. To quantify similarity, we calculated the Pearson correlation coefficients between samples and the expected composition (Figure 3B). Three of the four replicates showed a relatively strong correlation with the expected composition (∼0.8), while replicate P2 shows only a poor correlation (0.62). Upon closer inspection, four species (*Salmonella enterica, Staphylococcus aureus, Stenotrophomonas maltophilia* and *Pseudomonas fluorescens*) showed significantly higher abundance than the expected value of 4.26%. For *S. enterica* and *S. aureus*, the increased abundance is expected as multiple strains from the same species were combined. For the S. aureus strains, more than 80% of peptides (as listed in the protein fasta file) are shared between them. The S. enterica strains were only represented with a single species-level proteome. Furthermore, *P. fluorescens* was strongly overrepresented in samples P2 (17.8%) and P4 (9.8%), while being close to the expected fraction (4.3%) in samples P1 and P3. The overrepresentation was observed regardless of whether quantification was based on peptide or protein counts, and independent of whether counts or signal areas were used. Moreover, all runs showed comparable peptide intensity profiles, indicating that the deviation from the expected profile was not due to the accumulation of difficult-to-elute peptides across runs (Figure S1). Therefore, P. fluorescens may be genuinely more abundant in replicates P2 and P4. On the other hand, a few species were systematically underrepresented across all samples. For example, *Rhizobium johnstonii*, and *Rhizobium leguminosarum* account both only for approximately 1% of the total composition, and the algae *Chlamydomonas reinhardtii* was only observed at approximately 1.5%, as opposed to the expected 4.3%. In addition, most phages were underrepresented (e.g. 0.1% compared to expected 0.41%), and one phage (*Escherichia phage M13*) was not detected at all. While non-unique peptides can bias the obtained composition, phages may be further underrepresented due to their small number of proteins (usually some 50–150 proteins), resulting in fewer spectral counts for the same amount of protein in a typical shotgun proteomics experiment. Four strains included in the provided protein sequence database (*Desulfovibrio vulgaris, Nitrosomonas europaea, Nitrosomonas ureae, Nitrospira multiformis*) were not part of the mock community, although a small number of peptide matches (<0.1%) were detected for these species. However, this low level of matches is consistent with the expected false positives remaining after applying the false discovery rate (<1%) to the database search outputs.

**Figure 3.**
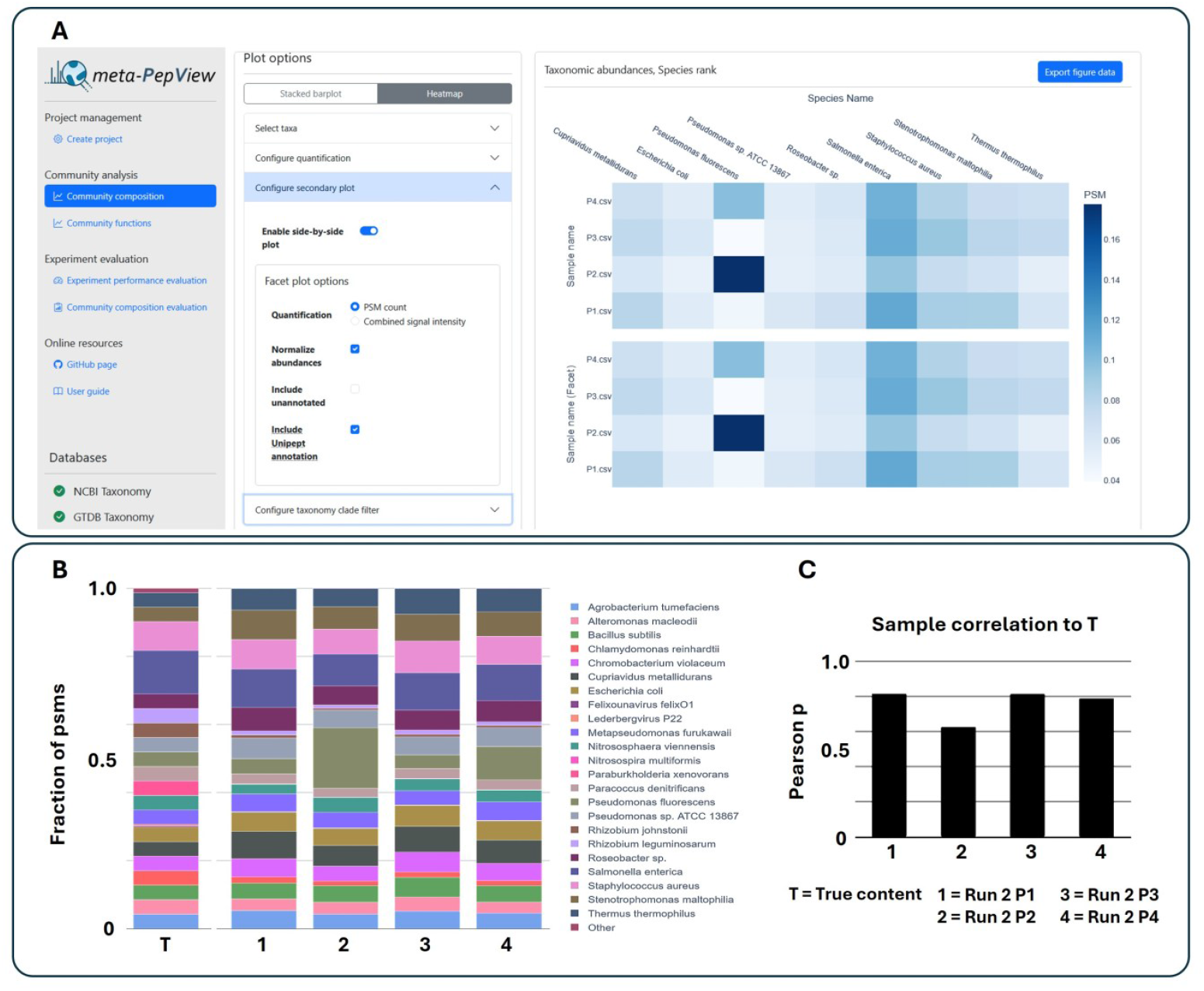
Taxonomic composition of Kleiner “equal protein community” replicates on species level, as shown by the meta-PepView community composition module. **(A)** Visualization of the community samples as heatmap in the “Community composition” module. **(B)** The observed species-level composition of four replicates (Run2_P1 to P4) from the Kleiner equal protein community (PXD006118) (76). Species composition was determined by meta-PepView counting all peptide spectrum matches assigned to each species. Peptides were identified via database searching using the provided mass spectrometric raw files and protein fasta file, with a 1% false discovery rate and a minimum of one unique peptide per protein. The expected composition as defined by Kleiner is shown by the left graph. **(C)** Pearson correlation coefficient (ρ) between the composition obtained by metaproteomics and the expected composition as defined by Kleiner and co-workers (76).

The accuracy of the abundance estimate of an organism within a community depends, among other factors, on the fraction of taxonomically unique peptides at a given taxonomic rank. To analyse for such deviations, the meta-PepView “Community Composition” module visualises the “taxonomic lineage drop-off”, i.e. the decrease in taxonomically unique peptides along the taxonomic lineage (Figure 4 A–C). For example, the drop-off curves for *S. maltophilia, R. leguminosarum* and *C. reinhardtii* are shown in Figure 4D. Interestingly, the total drop-off for *S. mallophilia* is only 6.2%, meaning that only a small fraction of peptides in this lineage are not uniquely assigned, while the average drop-off across all species in the community is 30.3%. On the other hand, *R. leguminosarum* shows a ∼90% drop-off across the taxonomic ranks, implying that only 10% of peptides were unique at the species rank. Interestingly, a major drop is observed at the species level, indicating high homology within the genus *Rhizobium* (*R. johnstonii*). Consequently, both species appear underrepresented when visualized strictly based on the LCA of each peptide, as implemented in meta-PepView. Another interesting case is the drop-off for *C. reinhardtii*, which is the only eukaryote present in the equal protein community, and therefore, nearly no drop-off is observed along its lineage. Nevertheless, *C. reinhardtii* is still underrepresented in the taxonomic composition graph (approximately 1.6% compared to the expected 4.3%).

**Figure 4.**
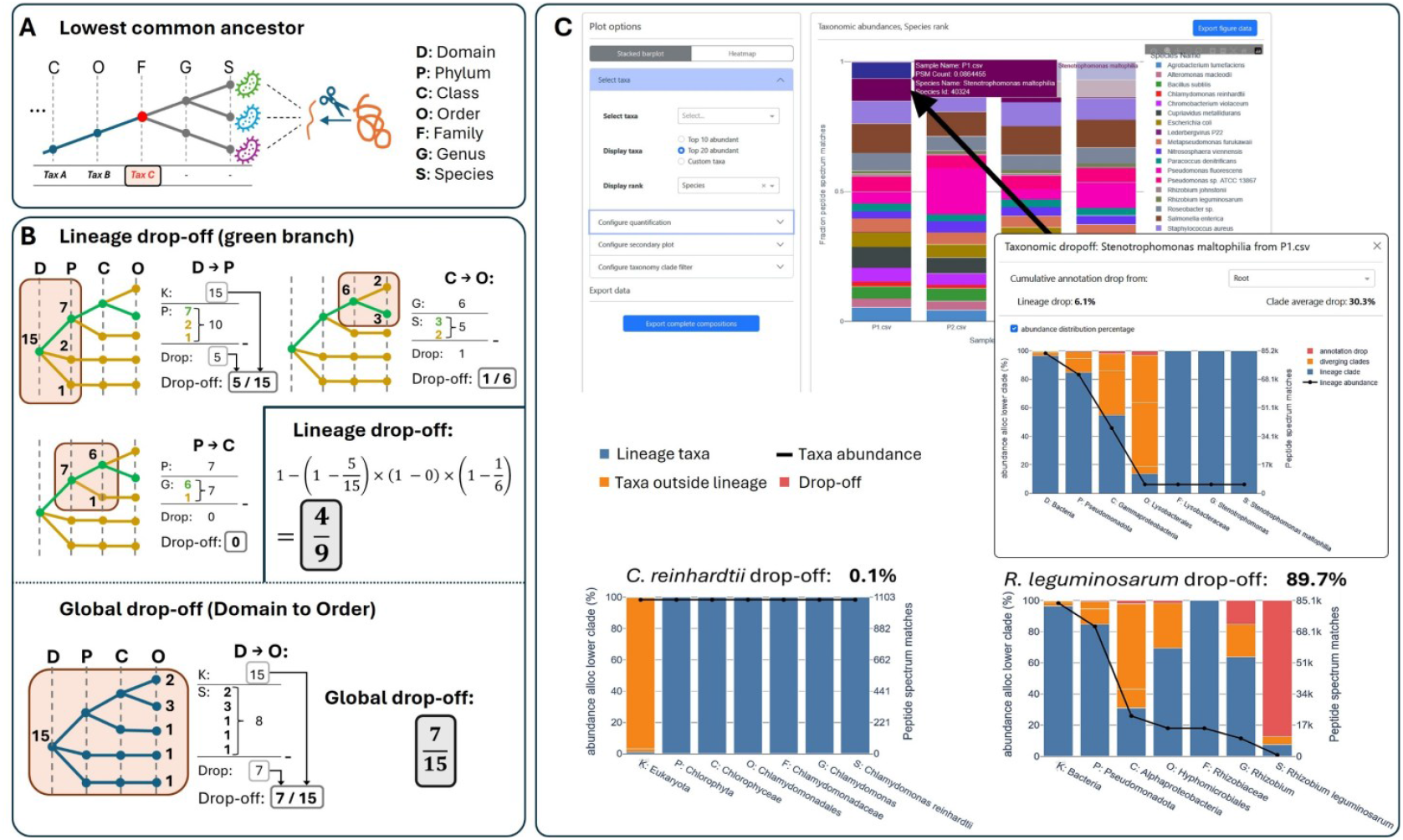
Taxonomic lineage drop-off and lowest common ancestor determination in meta-PepView. **(A)** Meta-PepView determines the Last Common Ancestor (LCA) for peptides based on taxonomy consensus from the provided reference sequence database. Each peptide is annotated with the taxonomic identifier of the LCA (“Tax C”), while lower ranks remain unannotated. **(B)** Meta-PepView calculates the cumulative drop-off for each organism across the entire lineage by determining, for each taxonomic level, the fraction of peptides that retain a valid annotation at the next lower rank (1 - drop-off). The product of these retained fractions gives the total retained fraction; and subtracting from 1 yields the cumulative drop-off. In contrast, the global (or average) community drop-off is calculated directly by dividing the number of peptides not annotated at the lowest rank by the number of peptides annotated at the root level. **(C)** The calculated drop-off rates for three species from the Kleiner equal-protein community. The drop-off graph is accessed by clicking the bar of the organism of interest. *S. maltophilia* exhibits a relatively low lineage drop-off (6.1%) compared to the total community drop-off (30.3%). In contrast, *R. leguminosarum* shows a very high lineage drop-off (89.7%), while *C. reinhardtii* displays virtually no drop-off (0.1%).

Since the community was assembled using equal amounts of protein extracted from cells, the presence of a rigid cell wall can be ruled out as a cause for the strong deviation from the expected value. However, *C. reinhardtii* is a phototrophic organism with an expected distinct functional profile. And indeed, the expressed functions show a strong enrichment of Chlorophyll binding proteins (Figure S2), where a few highly dominant proteins account for a large fraction of the protein biomass. This potentially biases quantification in shotgun proteomics, as their abundant peptides will be largely excluded from repeated fragmentation.

Next, we demonstrate meta-PepView by analysing a recently published SIHUMIx metaproteomics dataset (4). SIHUMIx is a simplified model of the human gut microbiome (83) that is widely used in meta-omics research. Spectral data from two samples (samples S05 and S08) as published by the CAMPI study (4) were database searched and *de novo* sequenced using PEAKS. Both sample S05 and S08 contained the same biomass material, but were prepared in different labs, employing different sample preparation methods and proteomic experimental conditions.

Here, we used two approaches for taxonomic annotation, a peptide-centric and a protein-centric approach. For the “peptide-centric” approach, as also used in the CAMPI study, samples were database searched with the tailored UniProt database provided by CAMPI article (4). Identified peptide sequences were furthermore annotated with Unipept (84), with organism filtering prior to LCA calculation to remove all non-SIHUMIx species. In the “protein-centric” approach, the SIHUMIx metagenome provided by the CAMPI study was used for database searching, with taxonomic classification performed using DIAMOND. An illustration of the obtained composition and function profile graphs for both samples is shown in Figure 5 (and SI EXCEL DOC). Both samples showed a very similar profile, with only a slightly higher abundance of *T. ramosa* in S05 compared to S08 (CAMPI: *E. ramosum*), which was also noted in the CAMPI study (4).

**Figure 5.**
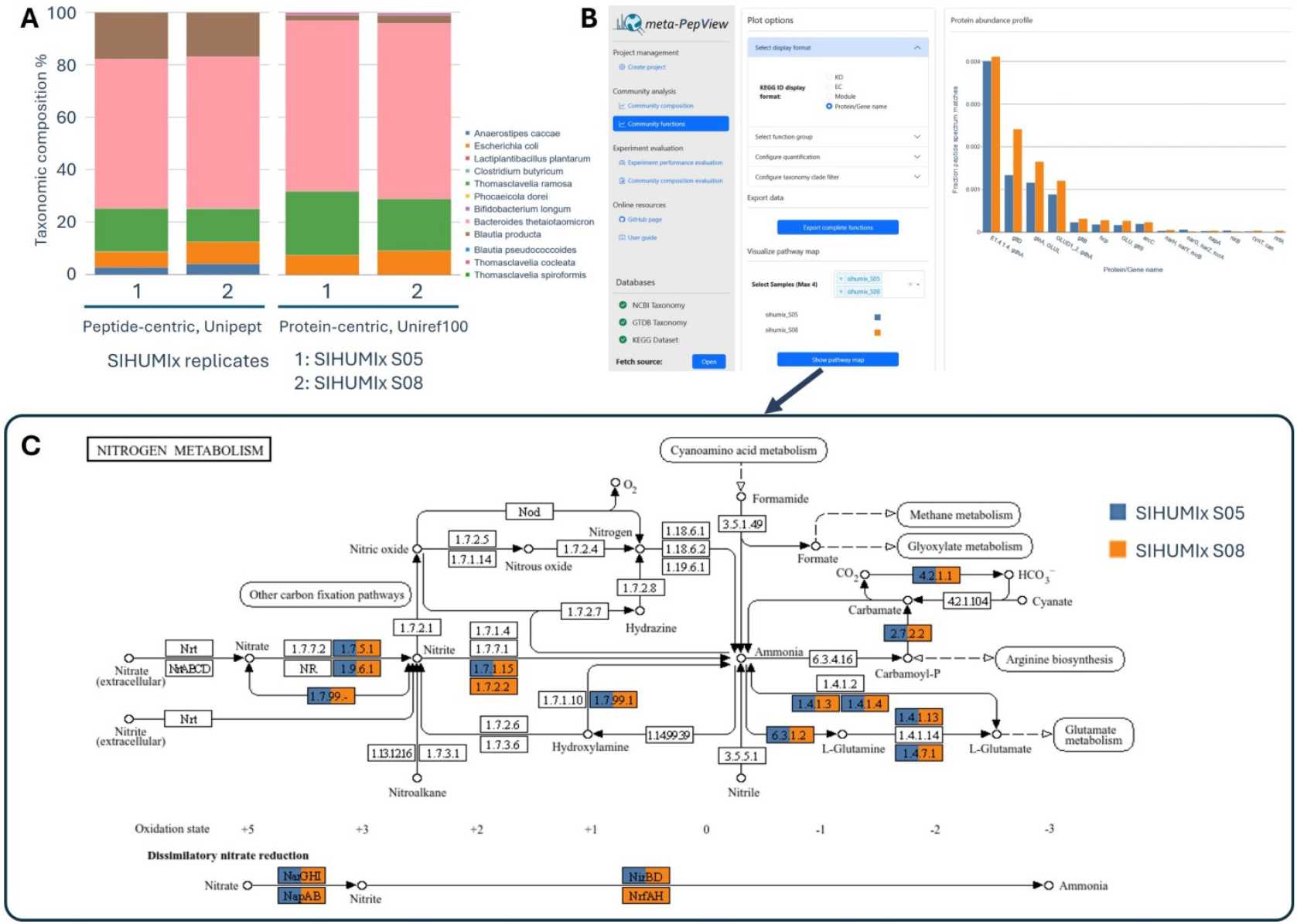
Example composition and function profiles for SIHUMIx samples as obtained by the meta-PepView “Community composition” and “Community function” modules. **(A)** The community composition of two SIHUMIx replicates (S05 and S08, as provided by the CAMPI study (4)) were displayed in meta-PepView as interactive bar graphs (shown as fraction of peptide-spectrum matches per species). Taxonomic annotation was performed “peptide-centric” and “protein-centric”. The “peptide-centric” approach was performed as described in the CAMPI study. Peptides were identified through database searching with PEAKS, using the UniProt-derived database as provided by the CAMPI study. Taxonomic annotation was done with Unipept (73), retaining only SIHUMIx taxa (4). For the “protein-centric” approach, peptides were database searched using the metagenome database provided by the CAMPI study, and proteins were taxonomically classified using DIAMON (80) and Uniref100. Both samples show a very comparable composition. However, the protein-centric annotation slightly differs from the peptide-centric approach **(B)** Meta-PepView gene abundance profile for both samples, presented as the fraction of total peptide-spectrum matches (y-axis) per gene (x-axis). Samples S05 and S08 were database searched with PEAKS, using the SIHUMIx metagenome database from the CAMPI study (4). The metagenome was functionally annotated with eggNOG, using default parameters. The gene abundance bar graph shows that nearly 0.5% of the metaproteome is assigned to glutamate dehydrogenase, reflecting its key role in nitrogen and carbon homeostasis. **C)** Meta-PepView KEGG pathway map of the nitrogen metabolism (KEGG: map00910). Meta-PepView submits the identified KOs to KEGG and overlays the resulting pathway maps with color-coded KO IDs corresponding to each sample. Visualization of the nitrogen metabolism pathway further reveals a complete DNRA pathway, explained by the presence of facultative anaerobes like *E. coli*, which can respire nitrate under anaerobic conditions.

Furthermore, in the CAMPI study, the major species observed in the peptide-centric approach (for both samples) were *A. caccae, B. thetaiotaomicron, E. coli*, and *T. ramosa* (CAMPI: *E. ramosum*). Interestingly, in our study we detected another abundant organism, *B. producta*, which accounted for more than 15% of the composition. The CAMPI study attributed the absence of *B. producta* to the reference database, as no reference genome was available in UniProtKB at that time. And indeed, in our case, the reference proteome for this organism (UP000289794) was already available.

In the protein-centric approach (using the metagenome for db searching) *B. producta* accounted for only 2– 3%, while A. caccae was detected only at trace levels. We therefore examined the metagenome database as a potential source of this underrepresentation using the meta-PepView “Community Composition Evaluation” module. This module compares the composition obtained from database search peptides, matched with the annotated metagenome proteins, to the composition obtained from taxonomic profiling using de novo peptide sequences, annotated with Unipept (26) (Figure S3, for sample S08). As also observed in the peptide-centric approach, the *de novo* peptide taxonomic profile shows a large fraction of *B. producta* and a notable amount of *A. caccae*. We then investigated the presence of *A. caccae* in the metagenome by matching *de novo* peptide sequences annotated to *A. caccae* against the metagenome database. This showed that only a small fraction of these sequences was covered by the metagenome database, and most were annotated to higher taxonomic ranks or to other organisms (e.g. for S08: only 89 out of 1107 peptides). This showed that the underrepresentation of *B. producta* and *A. caccae* in the protein-centric approach is due to insufficient coverage in the metagenomic database. This further highlights the influence of the reference sequence database on the observed composition (17, 27).

The “Community functions” module displays abundance profiles for protein function groups or visualizes expressed functions on KEGG pathway maps (Figures 4B and 4C). For example, the gene abundance profiles for the SIHUMIx samples shows that a significant fraction of the observed protein biomass (nearly 0.5% of the total metaproteome) has been assigned to glutamate dehydrogenase (GdhA) (Figure 4B). This enzyme catalyses an ATP independent route for the incorporation of ammonium into glutamate, which is essential for maintaining nitrogen and carbon homeostasis within the bacterial cell (85), and therefore a high enzymatic capacity can be expected. Interestingly, when investigating the nitrogen metabolism for the SIHUMIx sample, the dissimilatory nitrate reduction to ammonium (DNRA) pathway expression was found to be complete. Although the community is dominated by anaerobic fermenters such as *Bacteroides* and *Clostridia* (*Blautia* and *Thomasclavelia*), the presence of facultative anaerobes like *E. coli*, known to respire anaerobically using nitrate (86), supports this finding. DNRA converts nitrate to ammonium under anaerobic conditions, providing *E. coli* and other microbes with a strategy that can be energetically advantageous, while also supplying ammonium as a nitrogen source for the ecosystem.

### Example B: Meta-PepView spectral quality evaluation for SIHUMIx reference microbiome samples

In this section, we demonstrate the meta-PepView “Experiment performance evaluation” module with the SIHUMIx samples S05 and S08. Both datasets were generated from the same biomass starting material, albeit processed with different protocols and instrumental setups. The data were reanalyzed with PEAKS, the mzML spectral file, featureXML file, and both the database search psm and *de novo* peptide files were imported into meta-PepView. Using the experiment performance evaluation module, quality parameters related to chromatographic separation and spectral quality were extracted. For example, MS2 scans over time, displayed as the total ion current (TIC), were overlaid with histograms of chromatographic peak widths and high-confidence *de novo* peptide sequences (>90 ALC) over time (Figure 6A). It should be noted that S05 was analysed using a 460-minute gradient, while S08 used a shorter 240-minute gradient. That revealed that S08 shows a uniform distribution of high-confidence *de novo* sequences and peak widths, while S05 shows a steady decrease in high-confidence *de novo* sequences and increase in peak width over time. As a result, S08 resulted in a larger number of high-confidence *de novo* sequences despite the shorter run time. Furthermore, we accessed the spectral quality and resulting database searching scores. Lower spectral quality can arise when fragment-ion coverage is low or when fragmentation spectra are overly complex. Such issues may result from suboptimal isolation-window settings, poor MS2 ion transmission, or insufficient ion accumulation. This reduces database search scores and can also decrease the number of peptide identifications. For this, meta-PepView visualizes the distribution of high-confidence database search scores (Figure 6 B; top bars) alongside the fraction of high-confidence *de novo* sequences (>80 or >90 ALC) (Figure 6 B; bottom reversed bars). In addition, the fraction of “*de novo* only” (DN only) sequences are provided, which are spectra that provide high quality *de novo* sequences, but which are not matched during database searching. The absolute number of database search matches (Figure 6B; top bars) is larger in sample S05, which can be attributed to the longer run time.

**Figure 6.**
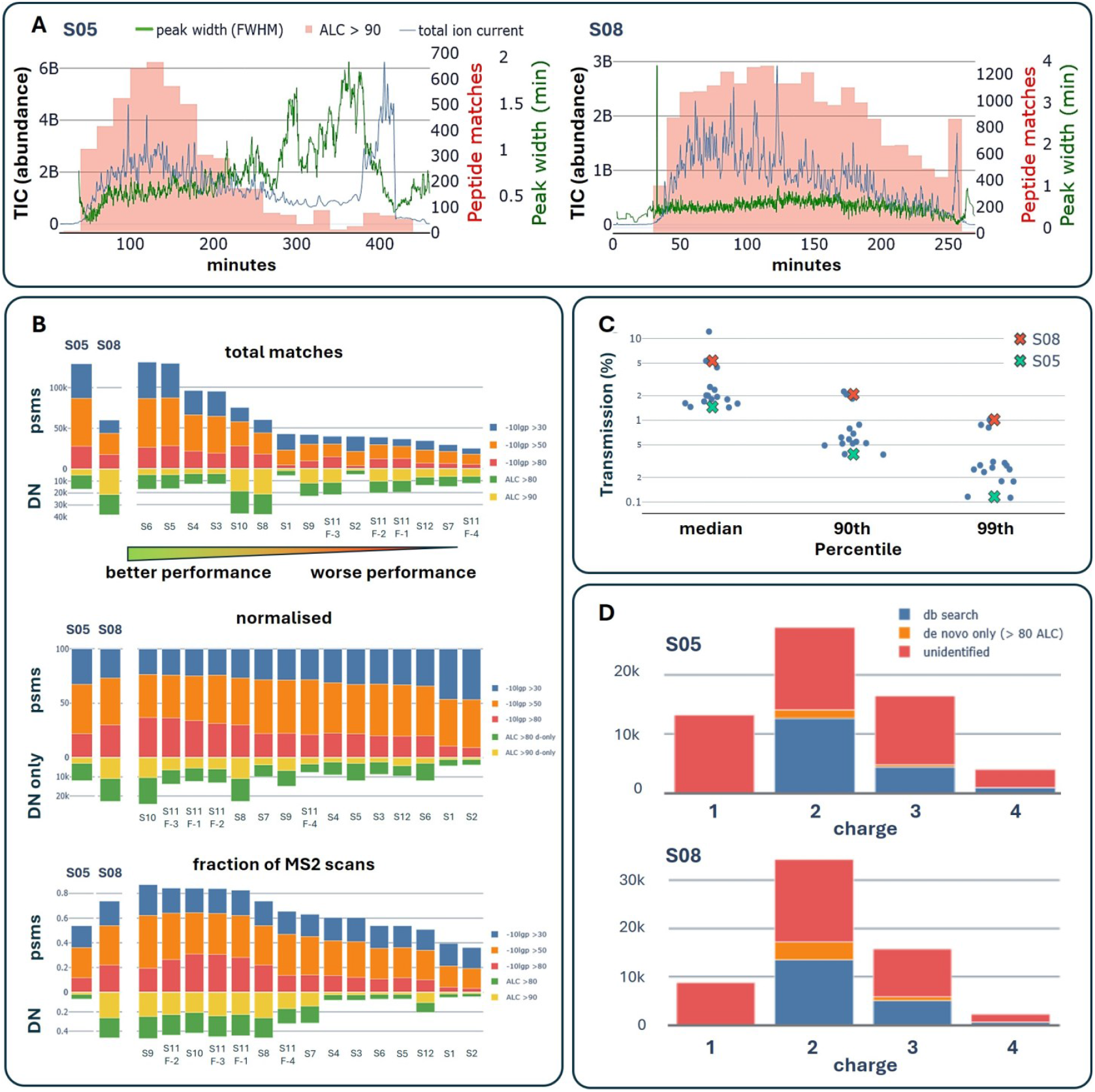
Meta-PepView chromatographic and spectral quality evaluation for SIHUMIx CAMPI datasets (4). **(A)** The two graphs show the ion current (TIC; left y-axis, blue line), high-confidence *de novo* sequence counts (ALC >90, right y-axis, red bars), and chromatographic feature peak widths (FWHM; right y-axis, green line) plotted over the run time (x-axis), for samples S05 (top graph) and S08 (bottom graph). **(B)** The graphs show the peptide identification performance for the SIHUMIx experiments S1–S12. S05 and S08 are shown on the left and labelled at the top of the bars, while the other samples are represented by the bars on the right, with sample numbers shown below the bars. The upper bars (“correct” y-axis) show the peptide spectrum matches grouped by confidence score (-10LgP, PEAKS), as total matches (upper graph), normalised to 100 (middle graph), or as fraction of the total MS2 scans (lower graph). The lower bars (reversed y-axis) show *de novo* sequences grouped by confidence score (ALC), as number of de novo sequences (upper graph), number of *de novo* only sequences (de novo sequences without database search match) (middle graph), and *de novo* sequence number as fraction of total MS2 scans (bottom graph). The datasets are ordered by number of database search matches (top graph), fraction of high-confidence database search matches (-10LgP >80) (middle graph), or by fraction size of database search matches (bottom graph). A large fraction of high-confidence database search matches and high quality *de novo* sequences implies high spectral quality, while a large fraction of de novo only sequences indicate a low reference database coverage. **(C)** The graph exemplifies the ion transmission efficiency for the SIHUMIx samples S05 and S08, benchmarked against the complete SIHUMIx dataset (S1–S12). The ion transmission efficiency is defined as: *TREFF = (MS*_*tic*_*/MS2*_*prec*_*) x (MS1*_*inj_time*_*/MS2*_*inj_time*_*), where MS1*_*prec*_ *is the precursor ion intensity, MS2*_*tic*_ *is the* summed ion intensity of the corresponding MS2 scan, *MS1*_*inj_time*_ *and MS2*_*inj_time*_ *are the scan injection times, respectively*. **(D)** Peak charge distribution of samples S05 (left graph) and S08 (right graph), shown as unidentified fraction (no database search or de novo identification), de novo only fraction (no database search identification), and database search fraction. A low fraction (and total abundance) of charge 1 features is desired, as these may indicate contaminations.

However, the database search psm score distribution (Figure 6 B; middle bars) shows a significantly larger fraction of high-confidence database search psms for sample S08. Moreover, S08 showed a substantially higher fraction of matched MS2 scans compared to S05 (0.7 versus 0.5, Figure 6B bottom bars), indicating a higher MS2 scan efficiency. For comparative purpose, Figure 6B also shows the complete SIHUMIx dataset as provided by the CAMPI study, which shows that the distribution of high-confidence peptide scores varies greatly across the entire SIHUMIx dataset. Interestingly, the samples with higher spectral quality (i.e. samples S10, S11) also provide a relatively large fraction of high-confidence DN only scores (Figure 6 B, middle graph). This indicates that the metagenomic reference sequence database may not fully cover the analysed community. This was also observed in the taxonomy composition evaluation, where the “*de novo* only” sequences annotated with the NovoBridge workflow (26) via Unipept provided a taxonomic profile that was biased towards certain microbes, including previously undetected ones such as *A. caccae* (Figure S3). Furthermore, the experiment performance evaluation module also allows to assess precursor ion transmission efficiency by evaluating the signal drop from MS1 precursor ions to their corresponding MS2 fragment ion intensities. The transmission efficiency for sample S05 aligns with the lower end of the performance range of the benchmark dataset, while sample S08 matches the best-performing reference data (Figure 6C). Finally, we investigated the presence of potential contaminants by visualizing the charge distribution of features (Figure 6D). Notably, a possible indicator for small molecule contaminants is the increased fraction of singly charged features. On the other hand, a large abundance of high charge state features (e.g. >6) may indicate incomplete digestion and elution of larger protein fragments. Comparing sample S05 to S08, both show a similar charge state distribution profile, although S08 shows a smaller fraction of singly charged features, potentially indicating a cleaner peptide sample. However, it should be noted that the samples were measured on different machines, with different settings and likely varying amounts of proteolytic digest, making the datasets not fully comparable. In summary, S08 showed a higher spectral quality, leading to better database search psm scores and high-confidence *de novo* sequences, where unmatched *de novo* sequences aligned to organisms not covered by the provided database. Despite differences in spectral quality, the fraction of database search matches at acceptable false discovery rate (<1% FDR) remained high for most SIHUMIx samples. The relatively small metagenome-based reference database (just over 20K proteins) likely allowed matching of lower quality spectra, with taxonomic profiles that were largely comparable.

### Example C: Evaluation of reference database coverage using meta-PepView

The database choice for database search psm is critical and can strongly influence metaproteomic results and conclusions (31, 87, 88). However, these effects can also be more subtle, affecting only specific subpopulations as seen for the SIHUMIx sample analysed above. This underscores the need for tools that assess sequence coverage and the accuracy of taxonomic annotations. Therefore, meta-PepView contains the “Experimental evaluation” module which evaluate the experimental performance (i.e. separation and data acquisition) as well as reference sequence database coverage. In the following part, we demonstrate this evaluation for a recently published Clostridium enrichment dataset (27, 89). The microbial composition as obtained from database searching using a metagenomics constructed database is shown in Figure 7A. As discussed recently, *Clostridium* appeared to be only a minor component in the enrichment, while a sulfate-reducing anaerobe (*Desulfovibrio*) accounted for >60% (27). To assess potential biases in the obtained taxonomic composition, we used meta-PepView to analyse taxonomic lineage drop-off, spectral quality, and reference database sequence coverage.

**Figure 7.**
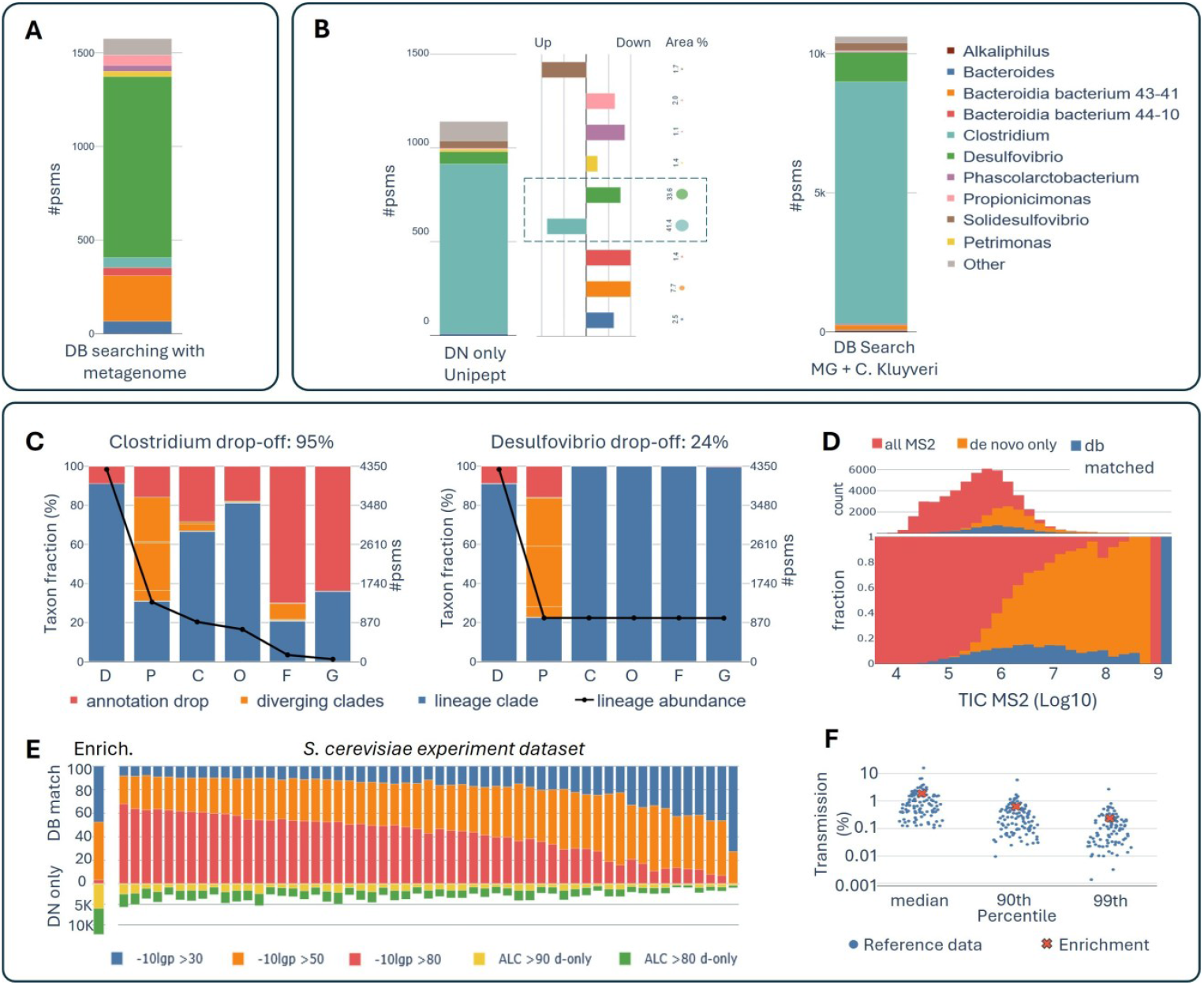
Evaluation of reference database coverage for Clostridium enrichment using meta-PepView. **(A)** The graph shows the genus-level composition as obtained from database searching using a metagenome-derived reference database. **(B)** The left bar graph shows the taxonomic composition of the Clostridium enrichment based on de novo sequences, using the NovoBridge workflow and Unipept database (>80 ALC). The adjacent panel displays the abundance differences between database-search (metagenome) composition and de novo profiling. The bar length represents the fold change of the de novo composition relative to the database-search (metagenome) composition, with *Clostridium* and *Desulfovibrio* highlighted. The circles indicate the area percentage of each microbe within the enrichment. The right bar graph shows the composition obtained from database searching using the metagenomic database supplemented with the *Clostridium kluyveri* reference proteome, which increased the psms for *Clostridium* from ∼1.5K to ∼10K. **(C)** The graph shows the taxonomic lineage drop-off for *Clostridium* (left graph) and *Desulfovibrio* (right graph). While *Desulfovibrio* shows a moderate lineage drop-off (∼24%), *Clostridium* exhibits a steep drop-off (∼95%) contributing to an underrepresentation at lower taxonomic ranks. **(D)** The upper panel shows the number of MS2 scans, while the lower panel shows the fraction of scans with a de novo identification or database search match as a function of MS2 intensity. Red bars show total MS2 scans, orange bars high-confidence *de novo* only sequences (>80 ALC) without a database match, and blue bars scans with peptide-spectrum matches. The data show that many high quality *de novo* sequences lack corresponding database matches, particularly for high-intensity peptide peaks. **(E)** The score distribution of database search peptide spectrum matches (from low to high: blue, orange and red) and fraction of high-confidence *de novo* only sequences benchmarked against a yeast reference dataset. The score distribution from the *Clostridium* enrichment (left bars) shows a low fraction of high-confidence database search matches compared to the benchmark dataset (red bar), while a relatively large fraction of spectra provided a high-confidence *de novo* sequence, but no corresponding peptide spectrum match (yellow and green bars), indicating that the reference sequence database does not fully cover the sequence diversity of the enrichment. **(F)** The plot shows the MS2 transmission efficiency, benchmarked against a yeast reference dataset, which confirms good transmission during analysis of the enrichment.

First, we investigated the taxonomic lineage drop-off for the expected dominant organism *Clostridium* and the apparently dominant organism *Desulfovibrio* (Figure 7C). While the *Desulfovibrio* lineage shows a 24% drop-off (i.e., 24% of the peptides were not uniquely assigned to *Desulfovibrio*), *Clostridium* exhibits a 95% lineage drop off. This means that only 5% of its peptides are uniquely annotated at the genus level, which results in an underestimation of *Clostridium* compared to other microbes. Furthermore, when checking the score distributions, the fraction of high-confidence *de novo* sequences (>80 ALC) compared to the total number of MS2 scans was very high, especially for peptides with high abundance (Figure 7D). Consistent with this, the fraction of psms was decreased for more abundant peaks. To contextualize the observed score distribution, the Clostridium data were benchmarked against a yeast reference dataset, that was measured over an extended period with the same instrumental setup (Figure 7E). This resulted in a relatively small fraction of strong database-search matches and many high-confidence de novo–only sequences, indicating that the metagenomic database did not fully capture the sequence diversity of the microbiome. Finally, the MS2 transmission efficiency was very good (Figure 7F), which means that even low-abundance peptide peaks should yield informative fragmentation spectra.

To further assess the reference sequence database coverage, we used the “Community composition evaluation” module to compare taxonomic profiles from database-search peptides (based on the metagenomic database) with profiles from high-confidence *de novo* sequences (>80 ALC) generated by the NovoBridge workflow and annotated using Unipept (42). The *de novo* taxonomic profile thereby confirmed the expected high enrichment for *Clostridium* (approximately 83%), while *Desulfovibrio* accounted only for 5.9% (Figure 7B, left graph). These were also the only two microbes with an area >5%. Finally, we supplemented the database with a *Clostridium kluyveri* proteome, as described by Kleikamp and co-workers recently (27). This increased the number of psms from 1.5K to over 10K, with Clostridium becoming the dominant microbe (Figure 7B, right graph). The absence of *Clostridium* in the metagenome was not further investigated but was likely caused by a community shift towards *Desulfovibrio* between sampling and sequencing (27).

## Conclusion

Here we introduce meta-PepView, a web-based modular platform for performance evaluation and interactive visualization of metaproteomic data, integrating major spectral, taxonomic, and functional annotation formats. The lightweight tool can be installed on any laptop or desktop via Docker or pip, runs in common web browsers, and requires only minimal field-specific expertise. The open-source code is freely available on the GitHub repository https://github.com/ramonzwaan/metapepview/. The growing importance of performance and quality evaluation in metaproteomics is addressed by evaluating chromatographic separation, spectral quality, spectral annotation, reference database coverage, and benchmarking against large sets of reference data. SIHUMIx and yeast reference datasets are provided with the installation. In addition, multiple datasets can be compared to support performance monitoring over time and to guide method development.

We demonstrate meta-PepView using two widely applied synthetic communities, the Kleiner equal-protein and SIHUMIx communities, additionally providing an extensive interpretation of these models for benchmarking metaproteomic experiments and data-processing pipelines. We further illustrate how small changes in taxonomic mapping or incomplete reference databases can alter experimental outcomes, underscoring the need to incorporate quality evaluation into metaproteomics. A detailed meta-PepView documentation is available to support installation and operation via https://ramonzwaan.github.io/metapepview/.

## Supporting information

SI DOC

## ACKNOWLEDGEMENTS

The authors thank all REThiNk consortium members for valuable discussions, Dita Heikens for support in the mass spectrometry facility, and colleagues from the Department of Biotechnology for their input. This work was supported by the Novo Nordisk Foundation grant NNF22OC0071498.

## CONFLICT OF INTEREST STATEMENT

The authors declare that they have no known competing financial interests or personal relationships that could have appeared to influence the work reported in this paper.

## Notes

### Competing Interest Statement

The authors have declared no competing interest.

