## Supplementary material for "Meta-PepView: a metaproteomics performance evaluation and visualization platform": SI DOC

### Table of contents

S1: Comparison of *P. fluorescens* peptide intensities between Kleiner equal protein replicates.

S2: Most abundant functions expression in *C. reinhardtii.*

S3: Comparison of taxonomic profiles between db search and de novo peptides for SIHUMIx S08.

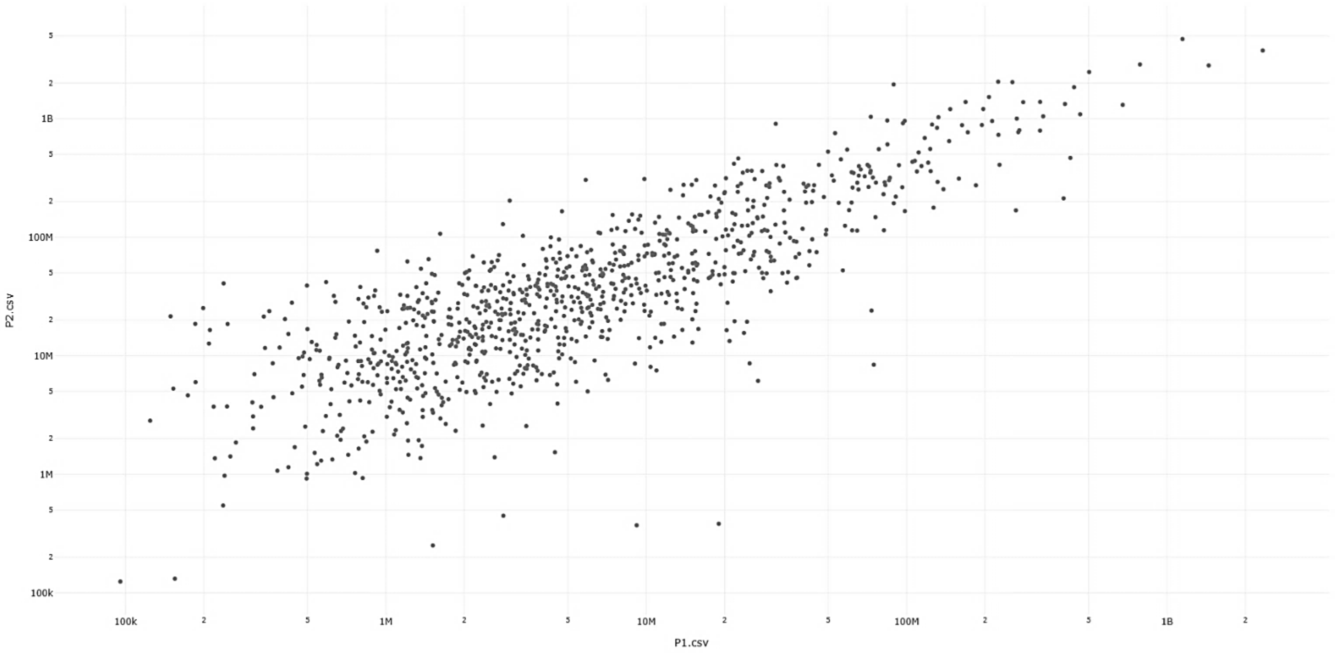


**Figure S1: Comparison of *P. fluorescens* peptide intensities between Kleiner equal protein replicates Run2 P1 and Run2 P2.** For each identified peptide sequence, intensities of sample P1 (x-axis) were plotted against P2 (y-axis). The plot shows a consistently higher abundance for *P. fluorescens* peptides in sample P2 over sample P1.

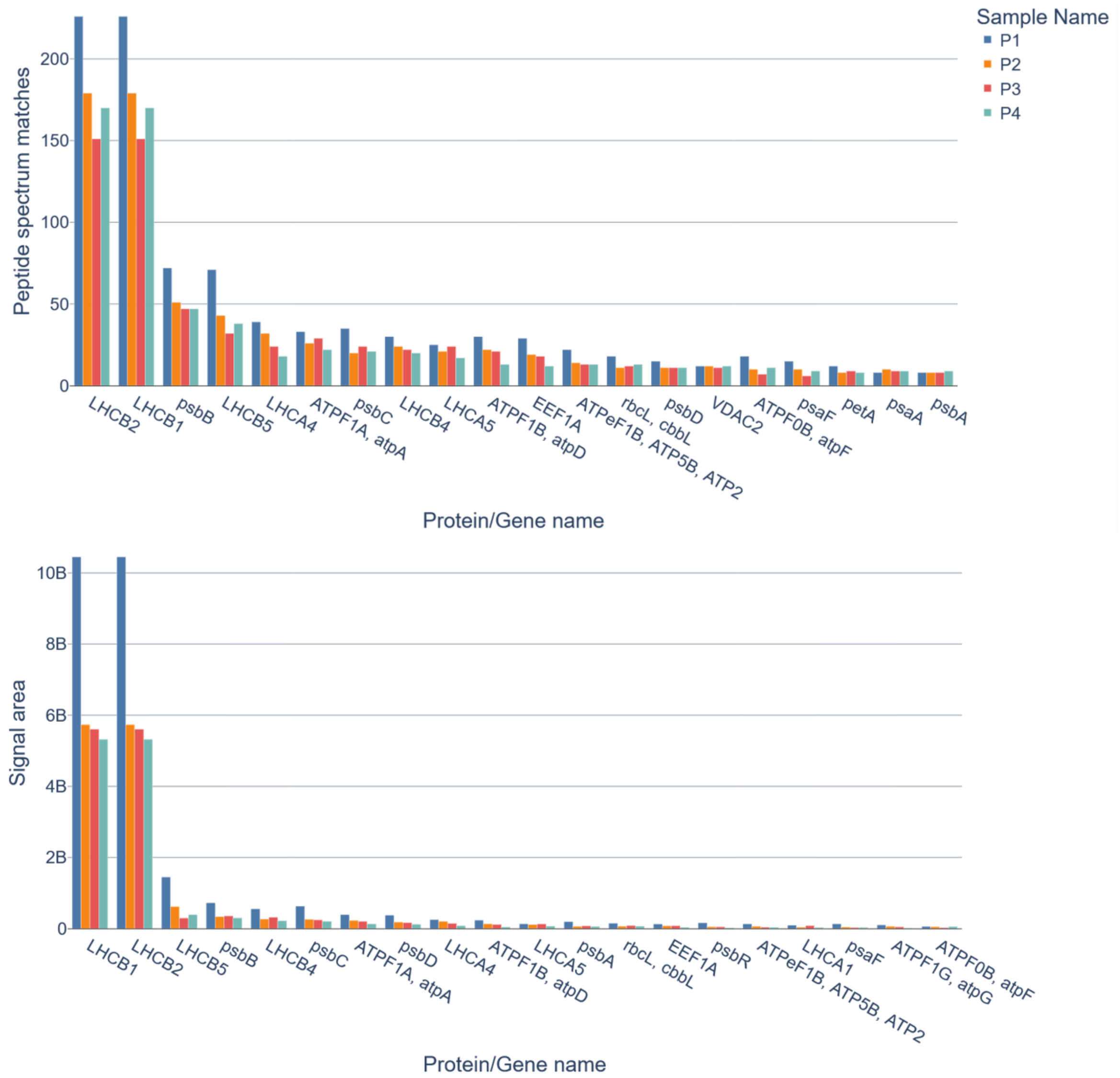


**Figure S2: Functional expression profile of proteins from *C. reinhardtii*.** The top 20 most abundant *C. reinhardtii* protein groups are visualized for each biological replicate (P1-P4). The top graph shows combined psm counts, while the bottom graph shows combined signal intensities. The graphs shows that the *C. reinhardtii* proteome is dominated by “light-harvesting complex” (LHC) proteins.

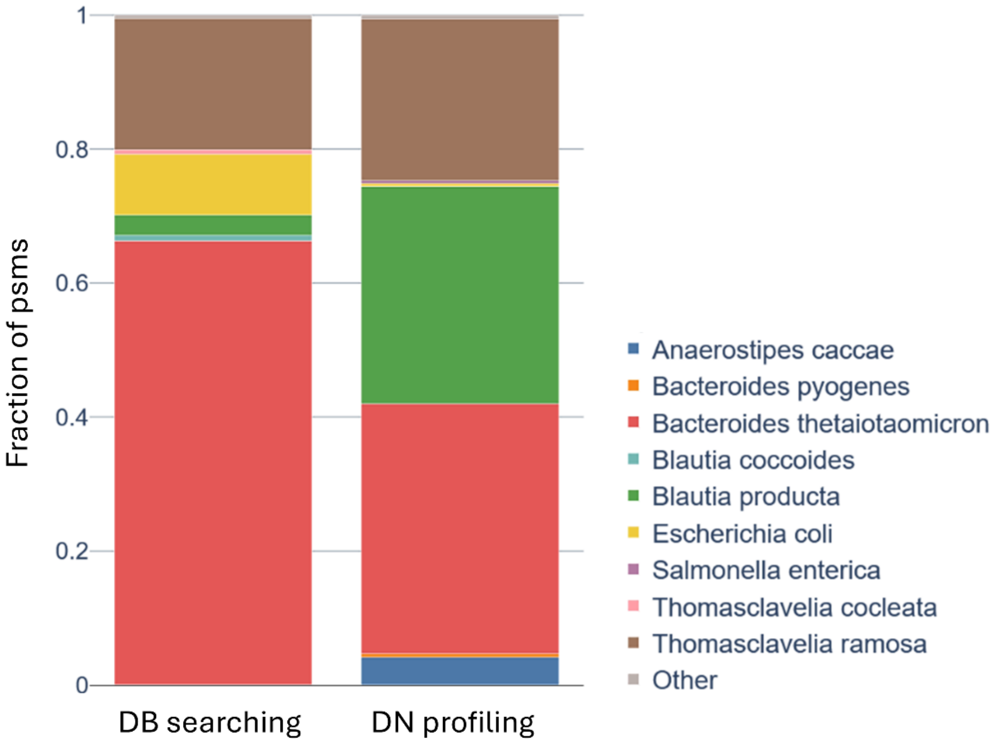


**Figure S3: Taxonomic composition obtained by db searching and de novo profiling for SIHUMIx S08.** "DB search" shows the taxonomic composition obtained from peptide matches using the SIHUMIx metagenome. "De novo taxonomy" shows the taxonomic composition obtained from de novo sequences that were taxonomically annotated by Unipept.
